# Predicting human mRNA isoform levels from site-specific splicing kinetics *in silico*

**DOI:** 10.64898/2026.06.16.732643

**Authors:** Zane R. Thornburg, You Jin Song, Jiaxi Yan, Kannanganattu V. Prasanth, Rohit Bhargava

## Abstract

Splicing of pre-mRNA can result in multiple possible mRNA isoforms per gene due to alternative splicing. The frequency at which individual isoforms occur depends on the intrinsic splicing kinetics of the pre-mRNA as well as intracellular chemical conditions. Computational modeling can potentially provide a platform to rapidly assess how variations in intracellular and environmental conditions, for example differential levels of regulatory splicing proteins, affect kinetics and resulting mRNA isoforms. Overcoming the vast combinatoric possibilities of splicing, however, has remained a significant challenge in modeling its kinetics. Here we report the development of a stochastic kinetic model of splicing that is extensible to most protein-coding genes in the human genome. Our model allows for variations in site-specific reaction rates as well as the ability to introduce additional splicing factors. We experimentally validate the predictive capability of our computational model by exploring the spliced isoform ratio of a target gene (*SRSF6*) under normoxia and hypoxia. This work provides a resource for quantitative, computational analysis of pre-mRNA splicing, allowing for a rapid computational-experimental approach to assess biological hypotheses.

**Author summary:** mRNA splicing is a key step in human gene expression with deep impact on the molecular processes determining cell physiology and affecting development and disease. Computational models of splicing are highly attractive to understand life processes but so far have been limited in directly accounting for the chemistry of splicing and are not extensible to most of the human genome. We report here a model that overcomes both of these challenges, providing a computational benchtop to probe splicing kinetics for most genes in the human genome. This model has the capability to rapidly pose biological hypotheses for experimental validation. As a targeted demonstration, we explore the effects of the pathologically-relevant chemical condition hypoxia on the pre-mRNA of a single gene.

## Introduction

Splicing is the process of removing non-coding RNA sequences (introns) and linking coding sequences (exons) in precursor messenger RNA (pre-mRNA). This mechanism is a fundamental process of eukaryotic life that enables genomes to code for more mRNA than its number of genes by linking exons in different combinations resulting in multiple mRNA isoforms per gene, each isoform serving a unique biological function. Since its discovery in 1977, [1, 2] splicing has proven to be a pillar of both our fundamental understanding of eukaryotic biology as well practical applications to improve human health. Dysregulated splicing is now understood to be a key feature of multiple molecular subtypes of cancer and is even the root cause of some genetic disorders [3, 4]. The success of drugs and therapeutics that modify splicing [4–7] motivate the need to more rapidly develop our quantitative understanding of the underlying biological, chemical, and physical mechanisms that control the outcomes of mRNA splicing.

While splicing leads to a diversity of mRNA that imparts great biological diversity, the same combinatoric complexity presents a challenge to understand and harness this phenomena for use. Computational modeling potentially presents a platform to probe this complexity of splicing as well as the large number of biochemical and biophysical variables affecting splicing kinetics. Simulations that incorporate biological processes that influence splicing outcomes can provide a workbench to test hypotheses before committing to experimental studies can significantly reduce experimental effort and provide support for mechanistic understanding. This has spurred the development of a significant number of machine learning and artificial intelligence models connecting sequence to predicted splice sites, possible mRNA isoforms, and even the ratio at which isoforms occur in different cell types or chemical conditions [8–15]. However, machine learning models do not provide direct insight into the influence of biological, chemical, and physical factors and dynamics that control splicing. Some mathematical and reaction-diffusion models, in contrast, have made progress in describing aspects of the fundamental dynamics of splicing [16–19]. One particular reaction-diffusion model made a prediction, for example, that spatial proximity of a gene to nuclear speckles affects the splicing outcome [17] - a hypothesis that has now been experimentally validated [20]. While the present mathematical and physical models can simulate realistic dynamics, none so far have been extensible to most of the human genome and frequently do not incorporate individual steps of the splicing cycle.

Here, we present a stochastic kinetic model of splicing that 1) is extensible to the majority of the human genome, and 2) allows for variations in chemical conditions and splice site-specific kinetics. The model leverages experimental measurements of the kinetic rates of splicing to predict not only which isoforms are formed, but also the frequencies at which they occur. The computational complexity to simulate the combinatorics of splicing was previously prohibitive. Recent advances in whole-cell modeling for bacteria [21] have pushed stochastic reaction software to simulate cell-scale reaction networks. Accounting for splicing of pre-mRNA that are still undergoing transcription is essential to mimic realistic splicing dynamics [22]. Recent advances in bacterial whole-cell modeling have also improved our ability in treating hybrid stochastic-deterministic reaction dynamics, allowing us to simultaneously simulate transcription elongation with splicing. Advances in quantitative measurements of the rates of individual reaction steps [23], improved resolution of individual splicing cycle states [24], and the availability of large quantitative transcriptomic datasets [25, 26] have increased the confidence in our ability to make accurate computational predictions.

## Materials and methods

### Overview of generalized kinetic model for mRNA splicing

To simulate the kinetics of mRNA splicing, we must first choose a set of reactions. We simplified the overall mechanism of splicing from the complete set of reactions [27, 28] to the mechanism shown in Fig 1A. Some known reactions are combined into ones that, for the purposes of determining splicing outcomes, act as kinetically equivalent lumped reactions summarizing the real mechanism. Additionally, we only explicitly treat the following spliceosome components in the reaction scheme: U1 small nuclear ribonucleoprotein particle (snRNP), U2 snRNP, and the U4+U5+U6 snRNP trimer. For an individual exon/intron pair, we assume that the U1 snRNP binds reversibly to the 5’ splice site of the corresponding pair [23]. The corresponding E complex can then undergo reversible pairing of the 5’ and 3’ splice sites before irreversible commitment to the A complex upon the ATP-dependent addition of the U2 snRNP [29]. Our model does not explicitly treat the branch point site-recognition binding of SF1 or the base pairing assistance of U2AF to the pre-mRNA and lumps the effects of site recognition into the U2 binding reaction. The U4+U5+U6 snRNP trimer then binds reversibly to the A complex to form the (pre) B complex. The (pre) B complex then undergoes an irreversible activation step in which the U1 leaves the complex to form an activated B complex. Typically, the stage described as the activated B complex lacks both U1 and U4, but because we do not explicitly simulate the assembly of the snRNP complexes, our activated B complex is functionally equivalent. Finally, we lump the remaining steps of the mechanism into a single splicing reaction removing the intron and linking the paired exons. In the model, the rates of each reaction arrow in the diagram have either been measured directly or were calculated from related experimental quantities (see Kinetic Parameters for Splicing Reactions).

**Fig 1.**
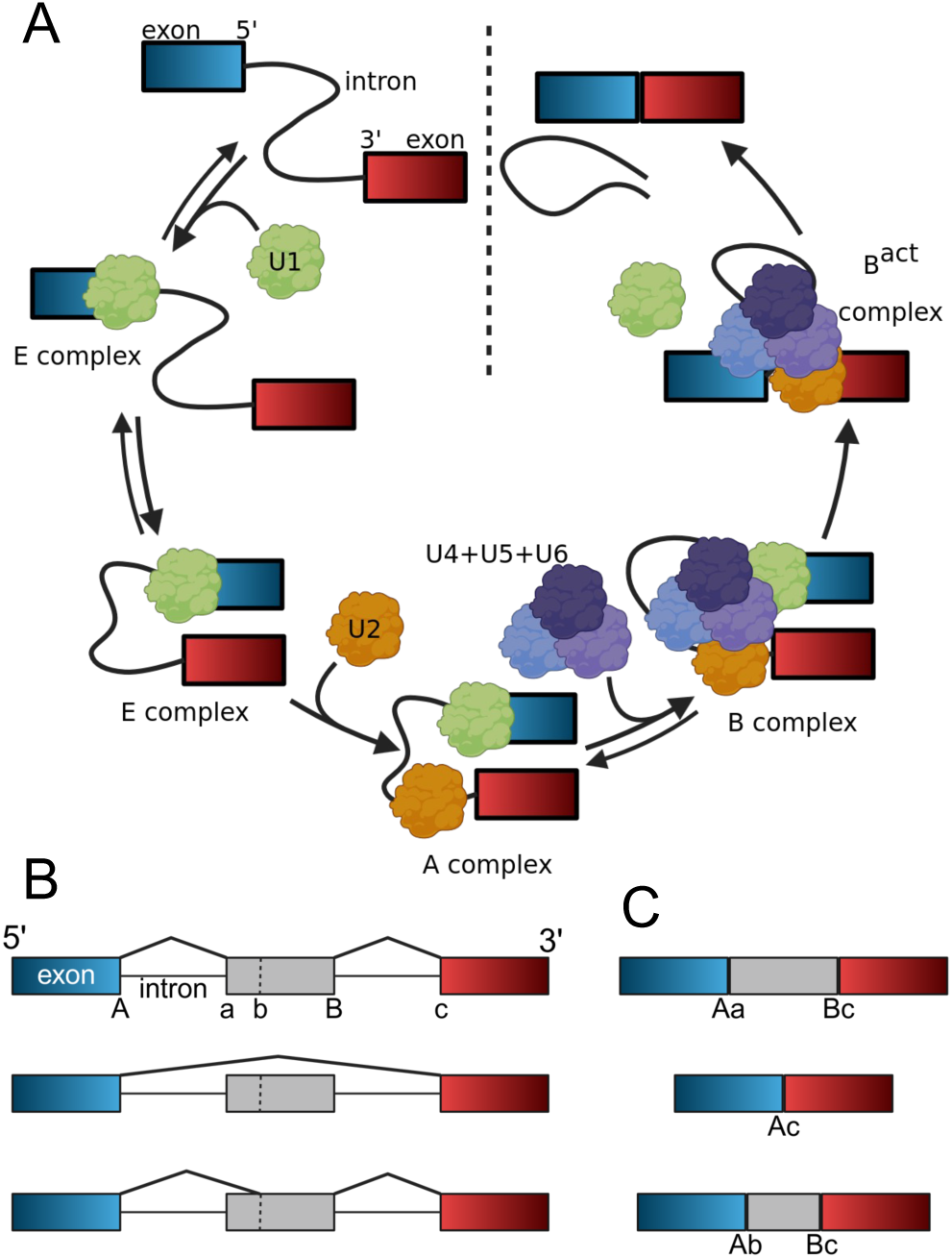
Overview of mRNA splicing reactions and outcomes. **(A)** Simplified splicing reaction cycle. **(B)** A single mRNA may consist of multiple possible splice site pairings. The model can predict removal of all introns and inclusion of all exons (top), alternative splicing through exon skipping (middle), and pairing with alternative splice sites, e.g. two 3’ splice sites for the same exon (bottom). **(C)** Splicing outcomes resulting from in the site pairings shown in (B). All panels created in BioRender. Thornburg, Z. (2026) https://BioRender.com/mb27moy and https://BioRender.com/j5uqm4j.

Mammalian gene pre-mRNA typically have more than one pair of exons; the average gene in the human genome has 9 exons [30]. Splicing in which all introns are removed and all exons are linked, as shown in Figures 1B and C, requires the reaction scheme in Fig 1A to be simulated for each splicing pair. The process of linking different combinations of exons is known as alternative splicing. Two forms of alternative splicing are: 1) exon skipping where splice site pairing occurs across an exon, resulting in it being removed from the pre-mRNA and not included in the mature mRNA and 2) alternative splice sites for the same exon resulting in different segments of the same exon being included in the mature mRNA. Both of these effects are shown schematically in Figures 1B and C. Therefore, not only do we need to model each neighboring splice site pair, but develop a model that can simulate arbitrary splice site pairing for any number and order of 5’ and 3’ splice sites in a pre-mRNA. The reduced complexity of our chosen reaction scheme made it possible to meet these criteria in a generalized reaction model for mRNA splicing where we treat each 5’ and 3’ splice site and their respective interactions independently.

There are two different times when splicing can occur in the lifetime of a pre-mRNA in our model: co-transcriptionally or post-transcriptionally. As the nascent RNA comes out of the RNA polymerase (Pol II) during transcription, individual splice sites become exposed sequentially. If spliceosome components are available to the exposed splice sites, the nascent mRNA can be spliced as transcription is ongoing [22]. In cases where the nascent mRNA is not exposed to spliceosome components during transcription or splicing reactions are slow for specific slice sites, splicing will not occur until the pre-mRNA is released into the nucleoplasm. Our model accounts for these dynamics in two ways. First, the transcription reactions can be set to produce the individual splice sites all at once (post-transcriptional splicing) or dynamically based on an input transcription speed and their position in the pre-mRNA sequence (co-transcriptional splicing). Second, in the case of the co-transcriptional reactions, the concentration of spliceosome components that the nascent-mRNA is exposed to is independently controllable from the total concentration of spliceosome components in the nucleus. Once the full pre-mRNA has been transcribed, the model switches the concentration of available spliceosome components to the total amount in the nucleus as the pre-mRNA is assumed to diffuse freely after transcription is completed (see Transcription elongation and custom procedures).

Gene expression is a partially stochastic process as it involves molecules that are low in concentration (nM - *µ*M) [31]. Splicing is no exception to this phenomenon, as exposed spliced sites are finite and cannot be treated as continuous concentrations, and the binding/unbinding of spliceosome components are random processes with likelihoods determined by their binding affinities. To simulate the stochastic nature of the splicing reactions, we use the chemical master equation solver implemented in Lattice Microbes [21, 32–34]. This method treats the reaction system, in this case the cell nucleus, as a well-stirred reaction vessel in which counts of individual molecules are updated using the Gillespie algorithm [33]. There are other methods that simulate stochastic reaction dynamics, but Lattice Microbes also provides the capability to simultaneously simulating deterministic processes [35], in the case of this model transcription elongation. The framework to run this model is Python-based and is publicly available on Github (https://github.com/Thornburg-Research/isosplicer/) and Zenodo (https://doi.org/10.5281/zenodo.20496770).

### Determination of mRNA isoforms from human genome annotation

The mRNA isoforms for the human genome were extracted from the NCBI entry GCF 000001405.38 by first downloading the Annotation Features file (GTF). The GTF file was then parsed into single-chromosome GTF files by a simple line search through the complete GTF file and copying all lines containing the following chromosome annotations into individual files: NC 000001.11, NC 000002.12, NC 000003.12, NC 000004.12, NC 000005.10, NC 000006.12, NC 000007.14, NC 000008.11, NC 000009.12, NC 000010.11, NC 000011.10, NC 000012.12, NC 000013.11, NC 000014.9, NC 000015.10, NC 000016.10, NC 000017.11, NC 000018.10, NC 000019.10, NC 000020.11, NC 000021.9, NC 000022.11, NC 000023.11, NC 000024.10. For each chromosome GTF file, the pyranges Python library was used to parse the file by first reading the list of “gene id” values to get the list of genes. For each gene, the GTF file was searched for all lines that were either an exon or gene “Feature” that were assigned the respective gene id and were annotated with a “Source” of BestRefSeq. The exon and gene data were then exported to single-gene excel files. The scripts to process the human GTF as described above are provided in the Github.

The user input for the model is a gene name that corresponds to the gene id’s from the human genome annotation and the folder containing the individual gene files described above. This input is used to read the single-gene excel file to determine the splice sites in the pre-mRNA and the possible isoform outcomes of splicing for the gene. First, the model reads the gene directionality. If the gene is on the froward (+) strand, all “Start” positions are mapped to the gene start site and 5’ splice sites and all “End” positions are mapped to the end of the gene and 3’ splice sites. The inverse of this mapping is done if the gene is on the reverse (−) strand. All splice site positions are then mapped based on the gene start position and are put in sequence order.

Once the splice sites are mapped to gene position, the model reads the list of “transcript id” that are in the gene annotation. For each annotated transcript, the list of exons included in the transcript is used to determine which splice sites were paired to form the transcript. The pairs in the transcript are then added to two lists: one list telling the model which splice site pairs form the transcript and a second that is a list of pairs that are allowed to form in the reaction model for the gene.

### Stochastic reaction solver

There are several programs available that can simulate stochastic reaction dynamics, and we chose to use Lattice Microbes [21, 32–34]. The choice of Lattice Microbes for a stochastic reaction solver is that it has the capability to interrupt the solver for user-defined custom procedures [21, 35]. Lattice Microbes calls this function a hookSimulation. The well-stirred chemical master equation stochastic reaction solver we use is the Gillespie Direct method (GillespieDSolver in Lattice Microbes). Directions for installation of Lattice Microbes including a file to create the required python environment are provided in the Github and Zenodo repositories.

### Kinetic parameters for splicing reactions

The kinetic parameters in the reaction scheme are either direct experimental measurements of the value or are calculated from indirect measurements. The full list is provided in Table 1. The U1 binding and unbinding rates come from direct measurements by single molecule spectroscopy [23]. Site pairing is calculated assuming a 2.2 min half life for a fully extended 100 nt intron [36] (using the extended intron length as the reaction radius) for a single pair of splice sites rounding to the nearest order of magnitude. Unpairing is assumed to match the the measured dissociation rate of U1-70K [37]. The U2 binding rate is calculated from a 0.15 min*^−^*^1^ reaction rate (5 minute reaction time) [29] assuming 1.2 *µ*M snrpB [38] in 40% HeLa cell extract [29] and rounding to the nearest order of magnitude. Binding of the U4+5+6 trimer is inferred from a measurement of 0.2 min*^−^*^1^ in yeast [39] with a 500 nm nucleus diameter and 100 trimers [40]. U4+5+6 unbinding is assumed to correspond to a slow dwell time of 1 min for U4 [41]. Activation is assumed to correspond to a short dwell time for U4 of ∼4 seconds [41]. The splice rate corresponds to a 15 second dwell time of U2 and U5 [42].

**Table 1.**
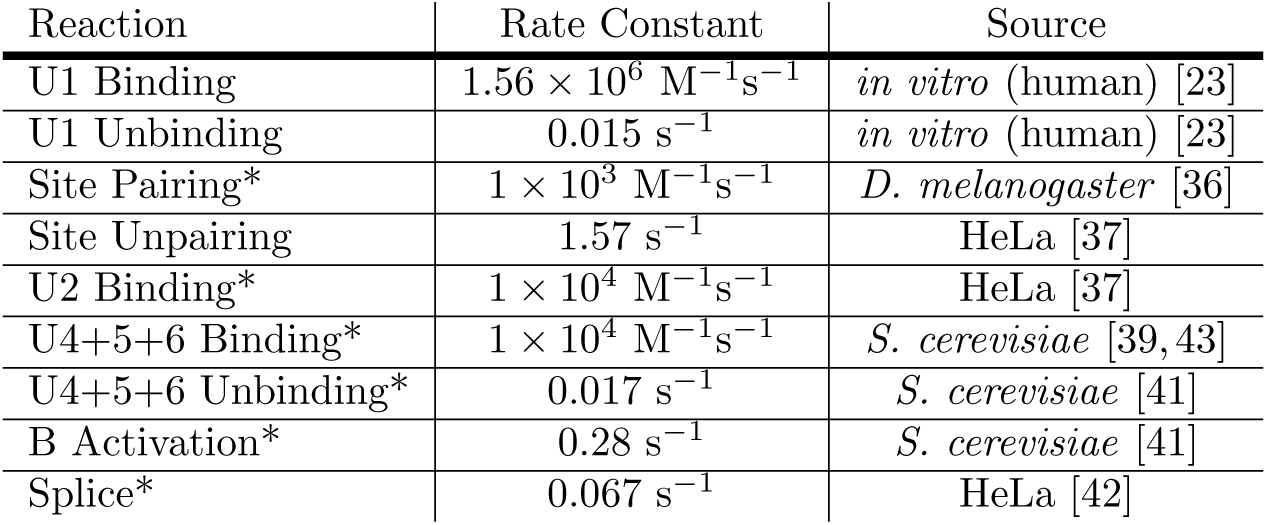
Reaction Rate Constants for mRNA Splicing and References for Their Experimental Measurements (*: Not directly measured, calculated from cited experiments)

| Reaction | Rate Constant | Source |
| --- | --- | --- |
| U1 Binding | $1.56 \times 10^6 \text{ M}^{-1}\text{s}^{-1}$ | *in vitro* (human) [23] |
| U1 Unbinding | $0.015 \text{ s}^{-1}$ | *in vitro* (human) [23] |
| Site Pairing* | $1 \times 10^3 \text{ M}^{-1}\text{s}^{-1}$ | *D. melanogaster* [36] |
| Site Unpairing | $1.57 \text{ s}^{-1}$ | HeLa [37] |
| U2 Binding* | $1 \times 10^4 \text{ M}^{-1}\text{s}^{-1}$ | HeLa [37] |
| U4+5+6 Binding* | $1 \times 10^4 \text{ M}^{-1}\text{s}^{-1}$ | *S. cerevisiae* [39, 43] |
| U4+5+6 Unbinding* | $0.017 \text{ s}^{-1}$ | *S. cerevisiae* [41] |
| B Activation* | $0.28 \text{ s}^{-1}$ | *S. cerevisiae* [41] |
| Splice* | $0.067 \text{ s}^{-1}$ | HeLa [42] |

### Initializing the reaction model for a single gene

Two sets of reactions are created for a gene upon model initialization: a transcription reaction and splicing reactions. For each splice site pair determined from the transcripts reported in the human genome annotation as described above and their respective splice sites, we initialize a set of chemical reactions following the scheme if Fig 1A and using the rate constants in Table 1. In the Lattice Microbes Python API, this involves defining a list of substrates, a list of products, and a rate constant per reaction. Substrates and products are spliceosome components, splice sites, and/or the intermediate splicing complexes shown in the mechanism.

Because the stochastic reaction solver deals with absolute numbers of molecules rather than concentrations, the model converts the concentration-dependent second-order rate constants into units of number by dividing by Avogadro’s number and a corresponding volume in which the reaction takes place. For the binding reactions of U1, U2, U4+5+6 trimer, and any additional factors, the volume is assumed to be the volume of the cell nucleus. Volumes of nuclei were calculated from experimental measurements of nucleus diameters as described below in initial conditions.

The reaction volume of the site pairing reaction is considerably smaller, as the splice sites are both part of a covalently linked molecule preventing them from drifting far apart. We chose to estimate the reaction volume for site pairing by using a physics-based theory [44] that estimates the hydrodynamic radius of an RNA segment as

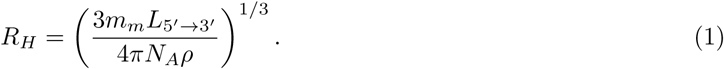

This method has been used previously for the purposes of estimating diffusion coefficients in a whole-cell model and we borrow its assumptions [32]: an average molar mass *m_m_* of 337 g mol*^−^*^1^ per nucleotide and a density *ρ* of 1.8 g cm*^−^*^1^. In this case, the length of the RNA segment *L*_5_*′_→_*_3_*′* is the distance in nucleotides from the 5’ splice site to the 3’ splice site.

### Transcription elongation and custom procedures

We use the hookSimulation functionality in Lattice Microbes to perform two functions every 1 second of biological time: final resolution of mRNA splicing and transcription elongation. The final products of the simulated splicing reactions are successfully spliced pairs of splice sites. To make the final product of splicing, a spliced mRNA, we implement a procedure to resolve splicing for the spliced site pairs. For each transcript from the genome annotation, we check each splice site pair required to make the transcript. If all site pairs for the transcript have been spliced, we remove the spliced site pair identifiers from the simulation and add one mRNA of the corresponding transcript.

If the simulation includes co-transcriptional splicing, the hookSimulation function also updates the transcriptional state of the nascent mRNA. The model assumes that the RNA polymerase (RNAP) starts at the gene start site. Once every biological second after transcription initiation, the model updates the position of the RNAP on the gene sequence by an amount corresponding to the input transcription elongation rate, default set to 2 kbp/min. Once the position of the RNAP is updated, the model checks the position relative to the list of splice sites. If the RNAP passed a splice site with its updated position, the splice site is added to the simulation to be available for splicing reactions. Once the RNAP reaches the end of the gene completing transcription, if the initial free fraction of spliceosome components and other factors was less than 1, the model sets the number of each respective particle to the total number in the initial conditions. This corresponds to the pre-mRNA having the freedom to explore the entire nucleus once transcription is complete.

### Intracellular initial conditions

There are two main components needed for the initial conditions of the model: the number of spliceosome components (and other molecules if included) in a single cell’s nucleus and the fraction of those molecules that are freely available to act upon nascent mRNA during co-transcriptional splicing. The number of U1, U2, and the U4+5+6 trimer in the nucleus were estimated using quantitative proteomics in HeLa cells [38]. The preoteomics study reports average single-cell protein concentrations. Representative components of each snRNP complex were selected to determine the whole-cell concentration of their respective snRNP: for U1 snrnp70 was reported to have a concentration of 1.66 *µ*M, U2 snrpb2 1.2 *µ*M, and U5 snrnp40 0.68 *µ*M. In the absence of cell line specific quantitative proteomics, we assume that the HeLa cell protein concentrations are transferable to other epithelial cell lines. We use estimated cell volumes for individual cell lines to determine the total number of splicing particles in the cell. MCF7 simulations assumed a cell diameter of 18 *µ*m and a nucleus diameter of 12 *µ*m [45]. Absolute counts of proteins are calculated using the proteomics concentrations from HeLa and the number of molecules per cell are determined using the cell diameter of the simulated cell type. For example, the concentrations for the proteins listed above correspond to 3.1 million U1, 2.2 million U2, and 1.3 million U4+5+6 per MCF7 cell. MCF10A simulations used a cell diameter of 20 *µ*m to calculate absolute counts of proteins per cell and used a nucleus diameter of 8 *µ*m (mode of reported distribution [46]). Additionally, to account for nuclear speckle localization of spliceosome components, we assume that only a fraction of the spliceosome components are available for co-transcriptional splicing. The fraction of U1 snRNP complexes that are in immobile states has been reported in the range of 70 to 80% [47, 48], so we assume that 25% of U1 snRNP are available for co-transcriptional splicing. We assume the same fraction for U2 and U4+5+6 complexes.

In the *SRSF6* normoxia/hypoxia simulations, the whole-cell concentration of SRSF4 from the HeLa proteomics was 0.98 *µ*M. Under the SRSF4 dispersal hypothesis, we assumed that 50% of SRSF4 were in an active phosphorylated state in both conditions. In normoxia we assumed that 10% of SRSF4 are available for co-transcriptional splicing and in hypoxia we assumed full dispersal of SRSF4 from speckles (100% available for co-transcriptional splicing). For the phosphorylation hypothesis, we assumed that 25% of SRSF4 were available for co-transcriptional splicing in both conditions (to match the fractions of available splicing components). In normoxia we assumed that 10% of SRSF4 are in an active phosphorylation state, and in hypoxia 90% of SRSF4 are in an active phosphorylation state.

### Simulating all genes with multiple isoforms

From the human genome annotation, a list of genes was compiled excluding genes with single reported transcripts and genes encoding micro-RNA. We iterated over this list, simulating 10 pre-mRNA per gene and running 30 simulations concurrently. Conditions were for a MCF7 nucleus size and spliceosome component abundances. Co-transcriptional splicing simulations assumed 100% of the spliceosome components in the nucleus were available for co-transcriptional splicing. The post-transcriptional splicing simulations assumed 0% of the spliceosome components were available to perform co-transcriptional splicing.

### Exploration of *SRSF6* Splicing Kinetics

We probed the reactions controlling exon 3 inclusion of *SRSF6* by adding SRSF4 binding reactions to the simulation. We assumed that the binding rate of U2 upstream of the “b” 3’ splice site (see Fig 3) was reduced to 0.001 M*^−^*^1^s*^−^*^1^ unless SRSF4 was bound downstream, in which case U2 binding is returned to normal levels. For SRSF4 binding downstream of the 3’ splice site for *SRSF6* exon 3 we assume a dissociation constant *K_D_* of 1 *µ*M: an intermediate binding rate of 1 × 10^5^ M*^−^*^1^s*^−^*^1^ and correspondingly an unbinding rate of 0.1 s*^−^*^1^. Inclusion of exon 3 was probed under 15 different sets of kinetic rates, one assuming uniform binding of all sites except for the 3’ site requiring SRSF4 and the other 14 each varying the rate of one or more reactions. The reactions that are changed all relate to either the binding of a spliceosome component or pairing rate for splice sites flanking *SRSF6* exon 3. All variations tested are listed in Table 2. All *SRSF6* replicates were simulated for 1320 seconds (biological time): 120 seconds for co-transcriptional splicing and 1200 seconds for post-transcriptional splicing.

**Table 2.** Kinetic parameters for exon 3 inclusion study in *SRSF6*.

| Parameter Set | Reaction | Site(s) | Rate Constant ($M^{-1}s^{-1}$) |
| --- | --- | --- | --- |
| 1 | Default | N/A | Table 1 |
| 2 | U1 Binding | C | $1 \times 10^5$ |
| 3 | U1 Binding | C | $1 \times 10^4$ |
| 4 | Site Pairing | B, b | $1 \times 10^2$ |
| 5 | Site Pairing | C, c | $1 \times 10^2$ |
| 6 | Site Pairing | B, b | $1 \times 10^2$ |
| | Site Pairing | C, c | $1 \times 10^2$ |
| 7 | Site Pairing | B, b | $1 \times 10^1$ |
| 8 | Site Pairing | C, c | $1 \times 10^1$ |
| 9 | Site Pairing | B, b | $1 \times 10^1$ |
| | Site Pairing | C, c | $1 \times 10^1$ |
| 10 | Site Pairing | B, c | $1 \times 10^4$ |
| 11 | Site Pairing | B, c | $1 \times 10^4$ |
| | Site Pairing | B, b | $1 \times 10^2$ |
| 12 | Site Pairing | B, c | $1 \times 10^4$ |
| | Site Pairing | C, c | $1 \times 10^2$ |
| 13 | Site Pairing | B, c | $1 \times 10^4$ |
| | Site Pairing | B, b | $1 \times 10^1$ |
| 14 | Site Pairing | B, c | $1 \times 10^4$ |
| | Site Pairing | C, c | $1 \times 10^1$ |
| 15 | Site Pairing | B, c | $1 \times 10^4$ |
| | Site Pairing | B, b | $1 \times 10^2$ |
| | Site Pairing | C, c | $1 \times 10^2$ |

### Cell lines

MCF7 cells were grown in RPMI 1640 medium supplemented with 5% fetal bovine serum and penicillin/streptomycin. Cells were maintained in a 5% CO_2_ incubator at 37*^◦^*C. For hypoxia treatment, cells were incubated in a 0.2% O_2_ and 5% CO_2_ hypoxia chamber for 24 hours. We confirm that the identity of all cell lines used in our study has been authenticated by STR profiling. All cell lines were regularly checked for mycoplasma using the ATCC Universal Mycoplasma Detection Kit.

### Immunofluorescence

MCF7 cells were seeded onto glass coverslips and cultured overnight until they reached approximately 40% confluency. Cells were then maintained under normoxia conditions or transferred to a hypoxia chamber and cultured at 0.2% oxygen for 24 h. Following treatment, cells were washed once with PBS and pre-extracted with 0.5% Triton X-100 in CSK buffer for 5 min on ice. Cells were then fixed with 4% paraformaldehyde in PBS for 10 min at room temperature and washed three times with PBS for 5 min per wash at room temperature on a shaker. Cells were blocked with 1% BSA in PBS at room temperature for 30 min. Primary and secondary antibodies were diluted in 1% BSA in PBS and incubated for 2 h and 30 min, respectively, at room temperature. Cells were washed three times with PBS for 5 min per wash after both primary and secondary antibody incubations. Nuclei were counterstained with DAPI at 1:5000 dilution in PBS for 5 min. Coverslips were mounted using VectaShield mounting medium. For imaging, z-stack images were taken with a 63x immersion objective using the ZEISS Axio Imager Z1 microscope with an AxioCam 506 Mono camera.

### Immunoblotting

Cells were collected by scraping and lysed in lysis buffer containing protease inhibitors and phosphatase inhibitors for 15 min on ice. Loading dye was added to the lysate and samples were then heated at 95°C for 5 min before loading onto and running on a polyacrylamide gel. After blocking in 5% milk, the membranes were incubated with primary antibodies, washed, then incubated with HRP conjugated secondary antibodies, and bands were developed with SuperSignal West Dura kit from ThermoFisher Scientific.

### Antibodies

Antibodies used in the present study include the anti-phosphoepitope SR proteins clone 1H4 (Cat# MABE50 from Sigma; IgG; WB 1:2000), SRSF4 (Cat# HPA050975; IgG; IF 1:99), SRRM2 (Cat# S4045 from Sigma; IgG; IF 1:1000), Tubulin (Cat# T9026 from Sigma; IgG; WB 1:20000), Goat anti-Rabbit Alexa Fluor 488 (Cat# A-11034 from Invitrogen), Goat anti-Mouse Alexa Fluor 568 (Cat# A-11031 from Invitrogen), HRP-conjugated Goat anti-Mouse IgG (H+L) Secondary Antibody (Cat# 115-035-003 from Jackson ImmunoResearch), and HRP-conjugated Goat anti-Rabbit IgG (H+L) Secondary Antibody (Cat# 111-035-003 from Jackson ImmunoResearch).

### Quantification of SRSF4 localization to nuclear speckles

Three biological replicates were analyzed for each condition, normoxia and hypoxia. For each biological replicate, four independent, randomly selected image fields were acquired and analyzed. Cells from both conditions were fixed, stained, and imaged in the same experimental session using identical staining conditions and microscope acquisition settings. After manual quality-control clean up, more than 40 cells were included for each condition.

For each image field, images were converted to 8-bit before analysis. Nuclei were segmented from the DAPI channel. Briefly, DAPI images were processed with background subtraction, followed by Gaussian blur. Nuclear masks were generated using Huang automatic thresholding, converted to binary masks, filled to remove internal holes, eroded twice to reduce peripheral halo signal, and subjected to watershed segmentation to separate touching nuclei. Nuclear regions were identified using the Analyze Particles function. Objects touching the image border were manually excluded.

SRRM2-posisitve nuclear speckles were segmented within each DAPI-defined nucleus mask. The SRRM2 channel was thresholded using Yen thresholding, converted to a binary mask, and processed with binary opening. SRRM2-positive regions within each nucleus were identified using Analyze Particles and combined into a single speckle ROI for each nucleus. The combined ROI was used as the unclear speckle mask.

SRSF4 intensity was measured within the DAPI-defined nucleus and the SRRM2-defined speckle ROI. SRRM2 nucleus integrated density was calculated as nucleus area times SRRM2 nucleus mean; SRRM2 speckle area fraction was calculated as SRRM2 speckle area divided by total nucleus area; SRSF4 nucleus mean was defined as mean SRSF4 signal in nucleus; SRSF4 nucleus integrated density was calculated as nucleus area times SRSF4 nucleus mean; SRSF4 speckle integrated density fraction was calculated as SRSF4 speckle integrated density divided by non-speckle integrated density; SRSF4 speckle enrichment was calculated as SRSF4 speckle mean divided by SRSF4 non-speckle mean; SRSF4 speckle mean was defined as mean SRSF4 signal in speckles.

Nuclei or speckles with unclear segmentation, merged nuclei, distorted morphology, or poor unclear boundaries, or ambiguous speckle masking were manually excluded from downstream analysis. Unpaired, non-parametric, Mann-Whitney test using GraphPad Prism 11. (ns p *>* 0.05; * p ≤ 0.05; ** p ≤ 0.01; *** p ≤ 0.001; **** p ≤ 0.0001)

### Processing of RNA sequencing data

The exon 3 inclusion levels for *SRSF6* were determined from published RNA sequencing data [26] using rMATS-turbo [49]. Levels were determined for MCF7 cells, each cell type having RNA sequencing reported for cells grown in both normoxia and hypoxia conditions. The previously published RNA sequencing dataset analyzed for MCF7 is deposited in NCBI GEO under GSE285077. Additionally, data was analyzed in the same manner for MCF10A cells prepared and sequenced using the protocols described in [26]. MCF10A cells were cultured at 37 degrees Celsius with 5% carbon dioxide and then grown in normoxic or hypoxic (0.2% oxygen) conditions for 24 hours before collection. RNA was extracted using Trizol and cleaned up using Qiagen RNeasy column. RNA sequencing data for MCF10A is deposited in NCBI GEO under GSE335740.

## Results and discussion

### Simulating splicing for the human genome

If we were to simulate splicing for every possible splice site pair, the number of possible mRNA isoforms would increase exponentially. For pre-mRNA that do not have alternative splice sites, the number of possible isoforms scales proportionally to 2*^N−^*^1^ where *N* is the number of introns. There are even more possibilities when alternative splice sites are present. Unfortunately, this makes prediction of an arbitrary number of isoforms per gene computationally impossible. To constrain our model, we only simulate isoforms, and consequently splice site pairings, that have been observed experimentally. We chose to constrain our model to the transcripts annotated in the GCF 000001405.38 NCBI RefSeq assembly (GRCh38.p12 Genome assembly). While there are newer annotations for the human genome, the RNA sequencing data we compare to later was mapped to this assembly and the genome pre-processing procedures we provide with the model will work as long as the data files provided by NCBI maintain the same structure and nomenclature. With these constraints shown in Fig 2A, we reduce the complexity of the simulations dramatically. For a genome-scale model, the maximum number of isoforms assuming all were possible would be 2^77^ (10^23^) isoforms for the *DMD* gene, where adding the constraint of only simulating experimentally observed isoforms reduces the maximum to 82 observed transcripts for the *CELF4* gene. From the genes in the annotation, 25,479 have reported transcripts, 12,025 were reported with more than one observed transcript, 11,515 only have a single reported transcript, and 1,939 are micro-RNA.

**Fig 2.**
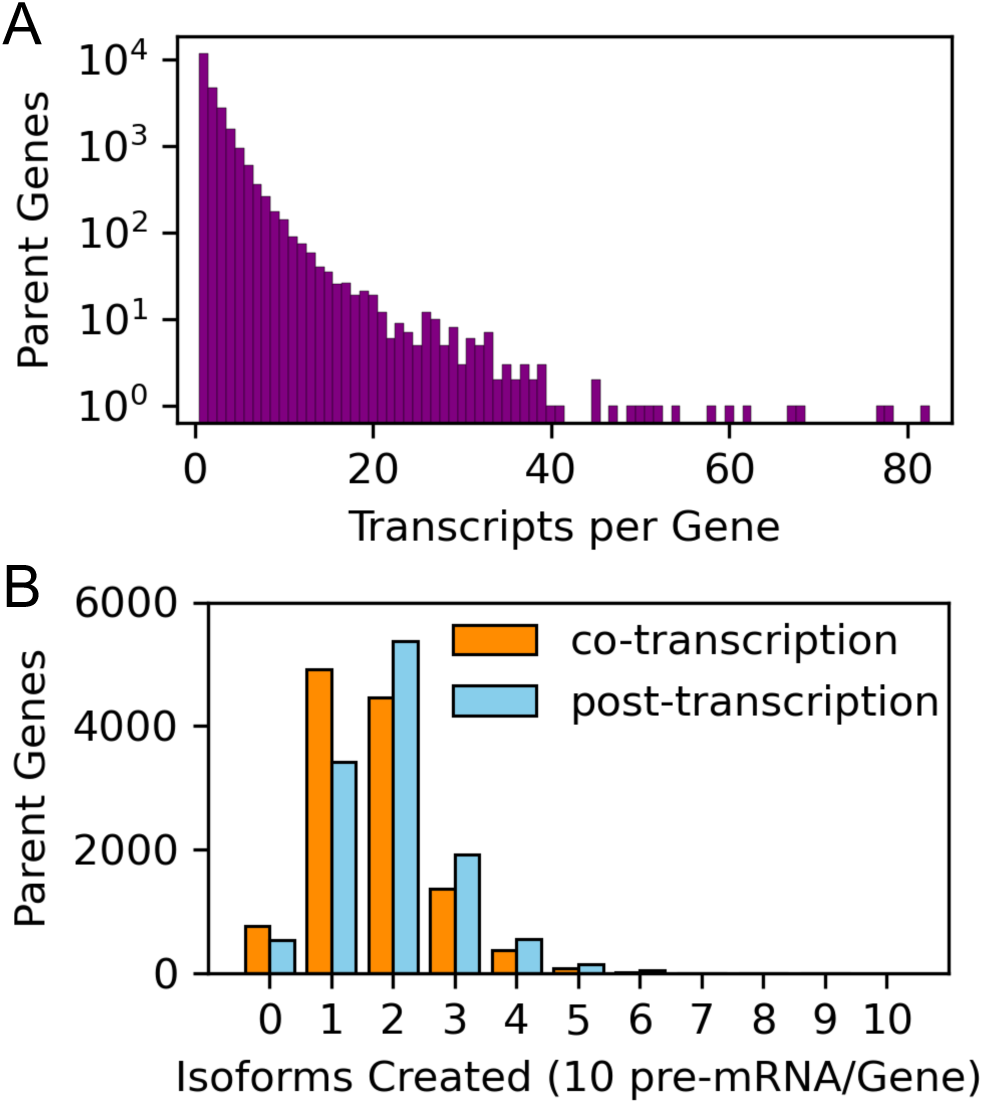
Genome-wide simulations of mRNA splicing. **(A)** The number of transcripts per gene reported in the human genome annotation GCF 000001405.38 that act as constraints for the splicing model. **(B)** For each gene with more than one reported transcript, we simulated splicing of 10 pre-mRNA in both co-and post-transcriptional splicing and counted the resulting isoforms. An isoform count of 0 indicates that no annotated isoforms were successfully spliced; most are due to incompatible mutually exclusive exons. Genes were simulated under conditions for an MCF7 cell nucleus.

To test the capabilities and limitations of our model, we simulated the splicing of 10 mRNA for each gene in the human genome annotation that was reported to have more than one possible isoform at two extremes: exclusively post-transcriptional splicing and co-transcriptional splicing with the nascent mRNA exposed to the entire pool of spliceosome components in the nucleus. We used a nucleus size for a representative MCF7 epithelial cell as the reaction volume [45] and corresponding concentrations of spliceosome components (see Methods). We show the number of isoforms that were successfully spliced out of 10 pre-mRNA per parent gene for each of the 12,025 genes under both conditions in Fig 2B. Generally we observe that exclusive post-transcriptional splicing results in more isoforms per gene that co-transcriptional splicing. This has also been predicted by other models [16] and is the expected result. In co-transcriptional splicing, a 5’/3’ splice site pair that is close in sequence will co-exist sooner than pairs that are distant in sequence, and the resulting temporal delays increase the likelihood that splice site pairs that are close in sequence will be spliced before distant splice sites are transcribed. No parent gene produced 10 mRNA isoforms among its 10 simulated pre-mRNA, but at least one parent gene resulted in each of 7, 8, and 9 isoforms.

The parent genes for which 0 mRNA isoforms were successfully spliced highlights some limitations of our model. The first limitation is that our model does not account for effects that truncate the ends of mRNA, specifically it does not account for multiple transcription start sites or termination sites and does not include post-transcriptional modification reactions beyond the splicing mechanism. Another limitation is some transcripts reported in the human genome annotation include intron retention. Our model currently recognizes these isoforms as incompletely spliced. Finally, a large portion of conflicts arise due to mutually exclusive exons. In the model, two pairs of splice sites that are observed in separate annotated transcripts, but not mutually in the same transcript, can both be spliced in the same pre-mRNA. If this occurs, the model will not recognize splicing as complete because the pre-mRNA does not result in any of the allowed mRNA isoforms.

For the model to be fully predictive, it should not only have the ability to simulate the formation of each isoform, but also be predictive of the ratio at which they occur in different cell types and intracellular chemical conditions. The genome-scale simulations described above assume that all splice sites and site pairs experience the same kinetics, which is not the case in reality. Each splice site and pairing are distinct due to their positions in their pre-mRNA sequence, affinities for spliceosome components, and interactions with other splicing factors. To enable exploration of the variations within a single mRNA, the model allows for user input of site-specific and pair-specific kinetics. We explore the isoform ratio more thoroughly for the gene encoding serine and arginine rich splicing factor 6 (*SRSF6*).

### mRNA isoforms of *SRSF6*

SRSF6 is itself a splicing regulator that is primarily known for playing a role in alternative splicing. The splicing of the pre-mRNA for *SRSF6* is an ideal candidate for probing the kinetic effects that control the ratio of isoforms. Only two transcripts have been annotated as possible outcomes of splicing as shown in Fig 3A: one isoform where its exon 3 is included and one where exon 3 is excluded. Exon 3 in the *SRSF6* pre-mRNA is a poison exon, a non-coding exon that when included adds a premature termination codon [50]. A binary outcome reduces the complexity of exploration of kinetics and validation of the model predictions.

**Fig 3.**
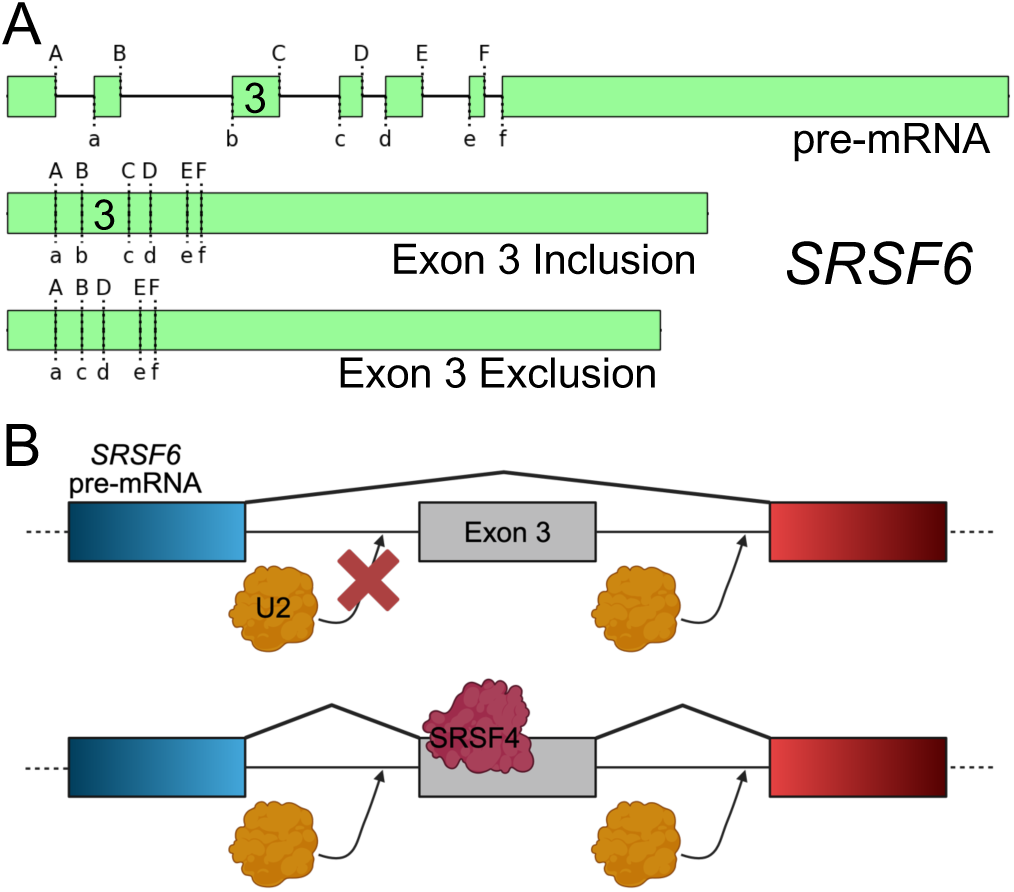
SRSF4 affects *SRSF6* splicing. **(A)***SRSF6* has two experimentally observed mRNA isoforms: one including and one excluding its third exon. **(B)** SRSF4 promotes inclusion of *SRSF6* exon 3 [51]. We simulate this by preventing the U2 binding step unless SRSF4 is present. (B) was created in BioRender. Thornburg, Z. (2026) https://BioRender.com/ou9nci5.

RNA sequencing (RNA-seq) data quantifying the isoform ratio has been reported in two different chemical conditions for MCF7 cells: normal oxygen (normoxia) and oxygen-depleted (hypoxia) [26]. Because the data consists of two chemical conditions, we have the chance to not only explore which splicing reactions control the isoform ratio of *SRSF6*, but also pose hypotheses about changes in intracellular conditions causing the observed isoform difference (17±1% exon 3 inclusion in normoxia, 39±2% in hypoxia).

### SRSF4 regulates splicing for *SRSF6* exon 3

It was previously reported that another splicing factor, SRSF4, promotes the inclusion of the *SRSF6* exon 3 [50, 51]. Typically promotion of exon inclusion of this kind involves binding of the splicing factor downstream of the 3’ splice site [52]. We identified a consensus SRSF4 binding sequence GAUGA [52] in exon 3 of *SRSF6* pre-mRNA 37 nt downstream of the “b” 3’ splice site in the reference genome (GCF 000001405.38). In the model, additional splicing factors can be added to the simulation and their association to individual splice sites can be controlled. The factor+site bound state then has its own binding rates for its respective snRNP (U1 or U2) that can be different from the site’s unbound kinetics. To give SRSF4 kinetic control of the inclusion of exon 3, we make an assumption that the binding of U2 upstream of exon 3 is extremely weak unless SRSF4 is bound downstream of the site as depicted in Fig 3B.

### SRSF4 alone does not explain the *SRSF6* isoform ratio

In determining the cause of the significant difference in exon 3 inclusion between normoxia and hypoxia conditions, one possible explanation is decreased nonsense-mediated decay [50]. However, the RNA-seq data indicates that the inclusion level difference observed for *SRSF6* is larger than the difference observed for poison exons of other *SRSF* pre-mRNA that are also known to be targeted by nonsense-mediated decay (22% for *SRSF6* compared to *<*10% for all others).

With our model we tested two hypothetical shifts in intracellular conditions that could impact *SRSF6* exon 3 inclusion. First, we hypothesized that the concentration profile of SRSF4 in the nucleus becomes more dispersed in hypoxia as shown in Fig 4A. A previous study reported a hypothetical mechanism where nuclear speckles dispersed in hypoxia in connection to increased *SRSF6* exon 3 inclusion [51]. As SRSF4 is known to concentrate itself in speckles [53], we proposed a hypothetical mechanism where SRSF4 disperses from speckles into the nucleoplasm in hypoxia, increasing the overall concentration of SRSF4 that *SRSF6* pre-mRNA are exposed to during co-transcriptional splicing.

**Fig 4.**
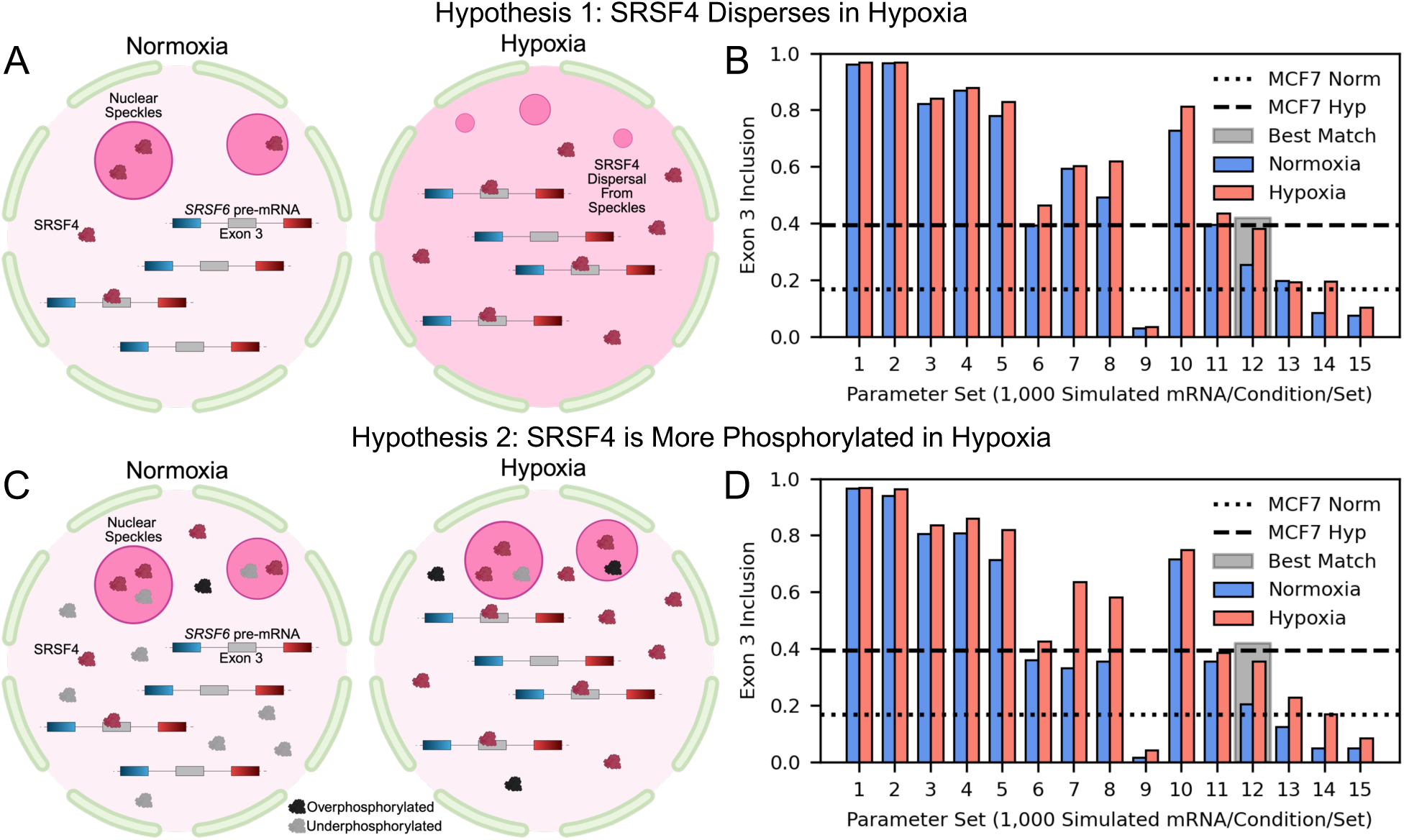
Two hypothetical mechanisms to control *SRSF6* splicing in normoxia and hypoxia. **(A)** Diagram showing a hypothetical mechanism of SRSF4 dispersing into the nucleoplasm from nuclear speckles in hypoxia. Dispersal of SRSF4 into the nucleoplasm results in more exposure for *SRSF6* pre-mRNA during co-transcriptional splicing. Created in BioRender. Thornburg, Z. (2026) https://BioRender.com/h88r5jt. **(B)** 1,000 mRNA were simulated per condition in 15 different parameter sets where site-specific reaction rate constants were varied from the uniform condition (parameter set 1). Exon 3 inclusion is calculated as the fraction of the 1,000 mRNA in which *SRSF6* exon 3 was spliced into the mRNA. **(C)** The second hypothetical mechanism to contrast normal and hypoxia conditions is a differential in the phosphorylation states of SRSF4. Red SRSF4 indicate a partially phosphorylated state that can bind to the *SRSF6* pre-mRNA with a high affinity. Grey and black SRSF4 indicate under- and overphosphorylated states, respectively. More SRSF4 are in active phosphorylated states (maroon proteins) in hypoxia than in normal conditions in this mechanism. Created in BioRender. Thornburg, Z. (2026) https://BioRender.com/1qykfs7. **(D)** 1,000 mRNA were simulated per condition using the same parameter sets in (B). Horizontal dashed/dotted lines indicate the experimental *SRSF6* exon 3 inclusion levels from RNA-seq [26]: 17±1% exon 3 inclusion in normoxia, 39±2% in hypoxia (n = 2 biological replicates).

A second explanation for the increased exon 3 inclusion is that more SRSF4 proteins are in active phosphorylated states in hypoxia. The phosphorylation state of SRSF4 is known to affect its ability to bind pre-mRNA and has been shown to affect the isoform ratios of other mRNA [54]. Phosphorylation of SRSF proteins is not a simple binary of phosphorylated or unphosphorylated, many phosphates can be bound to the RS domains of SRSF proteins, and it has been shown that only partially phosphorylated SRSF1 binds with a high affinity to its target RNA [55, 56]. We assume a similar effect applies to the binding of SRSF4 to its target RNA. Additionally, it was observed previously that hypoxia shifts the distribution of SRSF4 phosphorylation states to be more phosphorylated [57], likely due to upregulation of SR protein kinases (e.g. CLK1) by HIFs (hypoxia inducible transcription factors). Based on these observations, we assume that in hypoxia the vast majority of SRSF4 are in a phosphorylated state that can bind to the pre-mRNA and that the majority are underphosphorylated in normoxia conditions as depicted in Fig 4C.

For both hypotheses, we simulated the splicing of 1,000 pre-mRNA per condition assuming uniform splicing kinetics for each splice site pair. As can be seen in Fig 4B and D (parameter set 1 in both), these simulations predicted exon 3 inclusion in more than 90% of simulated mRNA. This indicates that SRSF4 alone is not enough to explain the experimentally observed isoform ratio, and that treating the kinetics for all splice sites and pairings uniformly is likely inaccurate for the *SRSF6* pre-mRNA.

### Site pairing rates control *SRSF6* splicing

To determine which reactions control *SRSF6* exon 3 inclusion, we varied the kinetics for the two 5’ splice sites (“B” and “C”) and the two 3’ splice sites (“b” and “c”) flanking exon 3 with 14 variations to the default kinetic parameters. 1,000 pre-mRNA were simulated per condition per parameter set; the complete list of parameter sets is in Table 2. We tried decreasing the affinity of the 5’ site C for U1 and the pairing rates for each respective splice site pair. Decreasing the affinity of U1 binding to C decreases the inclusion level of exon 3 (parameter sets 2 and 3 in Figures 4B and D), but not to the levels observed in RNA-seq. Further reduction of U1+C binding was not explored; we assume that it is unlikely for U1 to bind weakly to this 5’ splice site, as it has a close match (AU|GUAAGU) to the consensus sequence (AG|GU(A/G)AGU) [58, 59]. Also, our model suggests that this reaction does not result in an isoform level difference between normoxia and hypoxia. Of interest among other parameter sets that do not match the RNA-seq data, parameter set 7 (decreasing the B+b pairing rate by two orders of magnitude) predicts a significant amount of intron retention under both conditions in both hypotheses (roughly 20%, not shown in Fig 4).

We found that simultaneously increasing the pairing rate of B+c and decreasing the pairing rate of C+c by one order of magnitude each (parameter set 12) resulted in the closest match to the experimental RNA-seq data in the conditions of both hypotheses. The hypoxia inclusion levels are in decent agreement with the RNA-seq data, and the normoxia inclusion levels are slightly higher than the RNA-seq data. The higher inclusion in normoxia can be explained by our model not accounting for the fraction of mRNA that undergo nonsense-mediated decay. It is unclear what biological or chemical factors cause the varied site pairing rates required to obtain the experimental inclusion levels, one possibility could be RNA secondary structure of the nascent mRNA bringing the splice site pair that is distant in sequence (B and c) into close proximity in 3D space, effectively reducing the reaction volume and increasing the likelihood of pairing.

### Fluorescence imaging of SRSF4 dispersal in MCF7

While there is literature supporting the hypothesis that the phosphorylation state of SRSF4 increases in hypoxia, the hypothesis of dispersal of SRSF4 from nuclear speckles requires further validation. To experimentally determine if SRSF4 disperses during hypoxia in MCF7, we performed antibody staining and fluorescence imaging to characterize the SRSF4 distribution in MCF7 cells in normoxia and hypoxia. Representative nuclei are shown in Figures 5A and B. In the imaging we observed localization of SRSF4 to nuclear speckles, as well as other punctate condensates, in both normoxic and hypoxic cells. Speckle localization was determined by simultaneous imaging of SRRM2. We found that the total amount (integrated intensity, IntDen) of SRRM2 (Fig 5D) and the fraction of the nuclear area occupied by speckles (Fig 5E) were both reduced in hypoxia, indicating that SRRM2 is dispersing from speckles in hypoxia. The amount of dispersal we observe for MCF7 is less the the amount previously observed in HeLa [51]. We observe that although the total integrated density of SRSF4 in the nucleus remains the same between normoxia and hypoxia on average (Fig 5F), the intensity localized to speckles is reduced in hypoxia (Fig 5G). We confirmed that SRSF4 is enriched in speckles by determining the ratio of the average SRSF4 intensity in speckles to the average intensity in the the rest of the nucleus (Fig 5H), and we show that the concentration within speckles is similar between normoxia and hypoxia (Fig 5I). Our simulations indicate that significant dispersal of SRSF4 into the nucleoplasm in hypoxia could explain the experimentally observed difference in *SRSF6* exon 3 inclusion, however our imaging experiments indicate that some dispersal occurs in MCF7 but to a lesser extent than predicted by the simulations.

**Fig 5.**
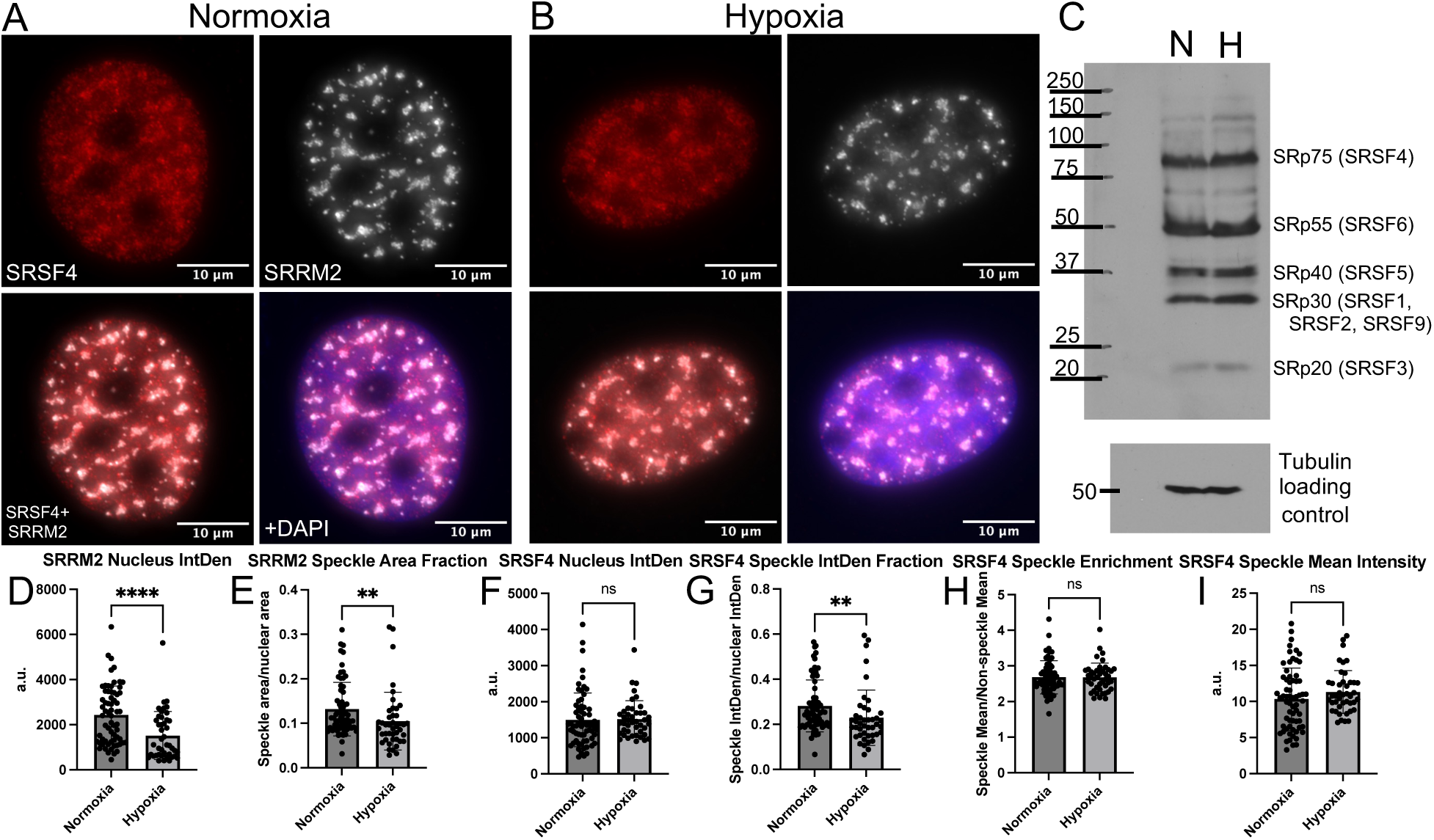
Experimental exploration of SRSF4 dispersal and phosphorylation. **(A)** Immunofluorescence imaging of SRSF4, SRRM2, and DAPI staining in a representative MCF7 nucleus under normoxia conditions. **(B)** Same color channels as (A), but under hypoxia conditions. **(C)** Immunoblotting of phosphorylated SRSF proteins under normoxia (N) and hypoxia (H). **(D)** Integrated (total) fluorescence intensity (IntDen) of SRRM2 in nuclei. **(E)** Fraction of the nucleus area occupied by speckles segmented from SRRM2 fluorescence intensity. **(F)** Integrated fluorescence intensity of SRSF4 in nuclei. **(G)** Fraction of total integrated SRSF4 intensity partitioned to speckles. **(H)** Ratio of SRSF4 intensity inside and outside speckles shows enrichment of SRSF4 in speckles. **(I)** Average SRSF4 fluorescence intensity in speckles. Scale bars for imaging are 10 *µ*m. Normoxia n = 65 nuclei; Hypoxia n = 43 nuclei. (ns p *>* 0.05; * p ≤ 0.05; ** p ≤ 0.01; *** p ≤ 0.001; **** p ≤ 0.0001)

The combination of previously reported experimental observations and our simulations indicates that an increase in the abundance of SRSF4 in active phosphorylation states is likely and can independently promote *SRSF6* exon 3 inclusion. The level of phosphorylated SRSF proteins was compared experimentally in MCF7 by immunoblotting using an antibody that preferentially recognizes the phosphorylated forms of all of the SRSFs as shown in Fig 5C. We observe only a slight increase in the total phosphorylated SRSF4 in hypoxia, but this does not tell us about the distribution of SRSF4 phosphorylation states [57]; this distribution has not yet been determined quantitatively. Because we observe some dispersal of SRSF4 from speckles in the fluorescence imaging, it is likely a combination of the two effects that cause the increase in *SRSF6* exon 3 inclusion. However, the small change we observe for SRSF4 dispersal indicates that the phosphorylation states likely shift more significantly. Or, given that the total SRSF4 appears similar between normoxia and hypoxia, if another effect could cause more SRSF4 to be available during co-transcriptional splicing in hypoxia than in normoxia conditions, that would offer another explanation.

### Testing transferability of kinetics to other cells

One of the most significant factors contributing to the predictive power of physics- and chemistry-based computational methods is the transferability of parameters between systems. For example, molecular dynamics has proven to be a powerful predictive tool, and its capabilities stem from the reliability in the atomic interactions parameterized in a force field (e.g. the bond strength of a carbon-carbon single bond) [60]. We tested the kinetic parameters that came closest to reproducing the RNA-seq data for MCF7 (parameter set 12, Fig 4D) under the phosphorylation-state hypothesis in a second system: MCF10A, non-tumorigenic mammary epithelial cells. To reflect the conditions of a different cell type, we recalculated the absolute numbers of spliceosome components and SRSF4 and changed the volume of the nucleus [46] (see Methods).

The exon 3 inclusion levels simulated in MCF10A cells shown in Fig 6 closely reflect the experimental inclusion levels. In normoxia conditions, the RNA-seq showed an inclusion level of 33±1% and our simulations predict 31.8%. Under hypoxia, the RNA-seq showed an inclusion level of 58±2% and the simulations predict 60.3%. This indicates that for related cells grown in similar conditions, kinetic rates can be transferable and predictive of isoform levels between cell types in our model.

**Fig 6.**
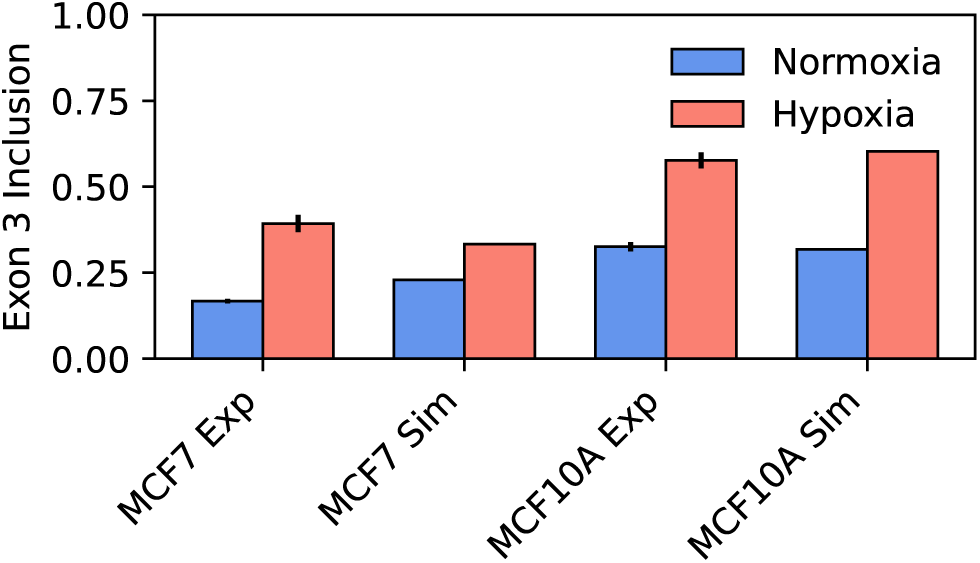
Transferring Kinetic Parameters to MCF10A. The kinetics from parameter set 12 in Fig 4D (MCF7) were transferred to conditions reflecting MCF10A cells. Simulated inclusion levels (Sim) are compared to experimental levels from RNA-seq (Exp). Each simulated parameter set and condition consists of 1,000 mRNA. Experimental inclusion levels include vertical error bars representing the standard deviation among biological replicates. MCF7 Exp: n = 2, MCF10A Exp: n = 2 biological replicates [26].

## Conclusion

Here, we presented a model of mRNA splicing that takes into account the reaction dynamics that drive splicing as well as the constraints and scale of the human genome. Our model can simulate splicing for most protein-coding genes in the genome annotation, and we showed that focused exploration of the kinetics for a single gene provides quantitative predictions for individual mRNA isoform levels. While it still has some limitations, for example lacking post-transcriptional modifications other than splicing, our model represents a leap forward in the ability to predict the outcomes of splicing through site-specific chemical reaction dynamics in a framework that is extensible to most of the human genome.

The ability to computationally probe the kinetic controls of splicing depends on our ability to both validate our predictions as well as improve on the model as new measurements are published. Quantification of the rates of individual reactions using single-molecule experiments will be increasingly valuable in improving the predictive power of models. Through our study of the kinetics of *SRSF6* splicing, we identified the progression from E complex to A complex, specifically splice site pairing, as an area where variations in reaction rates can strongly affect the outcomes of splicing. Identifying a range of realistic pairing rates will improve confidence in model predictions. Further measurements of binding rates for other splicing factors like SRSF4 and SRSF1 will help to computationally explore how variations in intracellular conditions affect alternative splicing.

## Data availability statement

All code to run the model is deposited on Github (https://github.com/Thornburg-Research/isosplicer/) and Zenodo (https://doi.org/10.5281/zenodo.20496770). The trajectories presented in the figures are also deposited on Zenodo and their corresponding parameter sets and analysis scripts to regenerate all graphs are deposited in both repositories. RNA sequencing data for MCF10A is deposited in NCBI GEO under GSE335740.

